# A binning tool to reconstruct viral haplotypes from assembled contigs

**DOI:** 10.1101/704288

**Authors:** Jiao Chen, Jiayu Shang, Jianrong Wang, Yanni Sun

## Abstract

**Motivation:** Infections by RNA viruses such as Influenza, HIV still pose a serious threat to human health despite extensive research on viral diseases. One challenge for producing effective prevention and treatment strategies is high intra-species genetic diversity. As different strains may have different biological properties, characterizing the genetic diversity is thus important to vaccine and drug design. Next-generation sequencing technology enables comprehensive characterization of both known and novel strains and has been widely adopted for sequencing viral populations. However, genome-scale reconstruction of haplotypes is still a challenging problem. In particular, haplotype assembly programs often produce contigs rather than full genomes. As a mutation in one gene can mask the phenotypic effects of a mutation at another locus, clustering these contigs into genome-scale haplotypes is still needed.

**Results:** We developed a contig binning tool, VirBin, which clusters contigs into different groups so that each group represents a haplotype. Commonly used features based on sequence composition and contig coverage cannot effectively distinguish viral haplotypes because of their high sequence similarity and heterogeneous sequencing coverage for RNA viruses. VirBin applied prototype-based clustering to cluster regions that are more likely to contain mutations specific to a haplotype. The tool was tested on multiple simulated sequencing data with different haplotype abundance distributions and contig sizes, and also on mock quasispecies sequencing data. The benchmark results with other contig binning tools demonstrated the superior sensitivity and precision of VirBin in contig binning for viral haplotype reconstruction.

**Availability:** https://github.com/chjiao/VirBin

## 1 Introduction

High genetic diversity within viral populations has been observed in patients with chronic infection with RNA viruses such as HIV, HCV, etc (Sullivan *et al.*, 2007; Perrin and Telenti, 1998). The genetic diversity could be caused by multiple infections of different strains or by mutations during the virus replication inside the host. In the latter case, the high replication rate, coupled with the low fidelity of the viral polymerase in most RNA viruses, results in a group of different but related strains infecting the same host, which is often termed as “quasispecies” (Nowak, 2006). Previous studies have revealed that patients with chronic virus infections, such as AIDS, are often the reservoir of new viral variants, which are likely produced during the replication process (MacLachlan and Dubovi, 2017). Because different strains could have very different biological properties such as virulence, transmissibility, antiviral drug resistance etc, characterizing the genetic diversity within viral populations is very important for developing effective prevention and treatment strategies. For example, if some strains have developed antiviral drug resistance, they may become the dominant strains and lead to treatment failure. Thus, characterization of the strain-level diversity of viral populations is indispensable for understanding the viruses and is of great clinical importance.

Sequencing viral quasispecies for genetic diversity analysis was one of the first applications of NGS (next-generation sequencing) technologies (McElroy *et al.*, 2014). Applying whole genome shot-gun sequencing to viral quasispecies does not require cultivation and can sequence divergent strains from known virus families. It thus has become a favored choice for characterizing the diversity of quasispecies.

Given the sequenced viral quasispecies, different types of analysis can be conducted to probe the genetic diversity. A relatively straightforward analysis is to understand the local diversity of known viruses by performing read mapping against reference genomes. While this type of analysis can produce a collection of local changes (mutations, insertions, or deletions) of the strains inside the quasispecies, it is not sufficient to infer the biological properties of the strains, which are more likely to be determined by multiple genes. In particular, epistatic interactions are abundant in RNA viruses, where the mutation of one gene masks the phenotypic effects of a mutation at another locus. Thus genome-scale reconstruction of the strains is essential for phenotype prediction of viruses (Töpfer *et al.*, 2014).

Reconstructing the genome-scale strain sequences inside quasispecies is often referred to as genome-scale haplotype reconstruction, where the genomes of strains are called haplotypes. The goal is to assemble short reads from sequenced viral populations into correct haplotype sequences. When the reference genome is available, read mapping can be conducted first to identify local mutations and then cluster the local mutations (or short contigs) into genome-scale haplotypes. When quality reference genomes are not available, which is often the case for emerging viruses such as SARS coronavirus, read mapping is not a very effective strategy to identify all mutations. Thus, *de novo* assembly is needed to stitch the reads into haplotypes.

With or without reference genomes, genome-scale haplotype reconstruction in quasispecies remains a computationally challenging problem. High similarity between haplotypes in the same quasispecies and the heterogeneous sequencing depth along the viral genomes present barriers to adoption of existing assembly programs. A recently published comparison showed that none of the tested haplotype reconstruction tools were able to successfully reconstruct the five known strains for a mock HIV quasispecies (Jayasundara *et al.*, 2014). We had the same observations when comparing several popular metagenomic assembly tools and haplotype assembly tools such as IDBA-UD (Peng *et al.*, 2012), IVA (Hunt *et al.*, 2015), SAVAGE (Baaijens *et al.*, 2017), MLEHaplo (Malhotra *et al.*, 2015) on the same data set (Chen *et al.*, 2018). Many methods output a set of contigs with various sizes that are much shorter than the genomes. With these outputted contigs from assembly programs, it still remains to infer the number of haplotypes and to match the contigs to their originating haplotypes. Thus, there is a need to cluster the contigs into different groups so that each group represents a haplotype. This step is called contig scaffolding or binning and has been applied for bacterial strain characterization.

Contig binning for viral quasispecies has its unique challenges. First, the goal of binning is to distinguish contigs from different viral strains rather than species. Thus, composition-based features such as tetranucleotide frequencies or GC contents are not informative enough to separate contigs from different haplotypes, which usually share high sequence similarity (over 90%). Tools that heavily rely on sequence composition-based features will not be able to estimate the number of haplotypes correctly. Second, RNA virus sequencing tends to be compounded by gene expression and fast degradation and thus the observed sequencing coverage along each haplotype, or even a contig, can be more heterogeneous than expected. In addition, if a contig contains a region that is common to multiple haplotypes, that region tends to have higher coverage than a haplotype-specific segment. All these challenges require carefully designed methods to use the coverage information for contig binning.

### 1.1 Related work

Although a number of contig binning algorithms have been developed (Wu *et al.*, 2014; Alneberg *et al.*, 2014; Kang *et al.*, 2015; Lu *et al.*, 2017; Quince *et al.*, 2017; Luo *et al.*, 2015; Truong *et al.*, 2017), they all possess limitations in distinguishing contigs from different viral strains of the same species. Most of the existing contig binning tools for microbiome sequencing data are designed for bacteria. These methods usually estimate the bin number by aligning metagenomic data to a pre-established marker gene database, and then assign assembled contigs to different bins using sequence composition information and read coverage levels. For example, MaxBin (Wu *et al.*, 2014) uses both tetranucleotide frequencies and contig coverage levels to assign assembled contigs into different bins.

Some binning tools (Lu *et al.*, 2017) leverage co-abundance of genes across multiple metagenomic samples. The rationale is that if two contigs are from the same bin, their coverage profiles across multiple samples should be highly correlated.

Recently, there are a couple of newly developed tools for strain level analysis from metagenomic data, such as Constrain (Luo *et al.*, 2015) and StrainPhlAn (Truong *et al.*, 2017). Both rely on species identification using clade-specific genes, then zoom in to identify the strains. However, both tools were mainly tested on bacteria.

Our method is designed to cluster contigs produced by existing assembly tools. There are another group of methods conducting haplotype reconstruction via read clustering (Ahn *et al.*, 2018; Barik *et al.*, 2018), which groups variant sites obtained by read mapping against reference genomes. These tools don’t usually output contigs and thus do not use contig binning.

Here we present VirBin, a method designed specifically for binning contigs derived from viral quasispecies data. The input to VirBin is a set of contigs derived from assembly tools. The output includes the estimated number of haplotypes, the grouped contigs for each haplotype, and the corresponding relative abundances. Unlike many bacterial contig binning tools that require multiple samples, our method works on a single sample.

## 2 Methods

The overall pipeline of our method is shown in Fig. 1. There are mainly two steps: (1) estimate the number of haplotypes by aligning contigs and identifying windows; (2) calculate relative abundances in each window and apply a clustering algorithm to group clusters of the same haplotype.

**Fig. 1.**
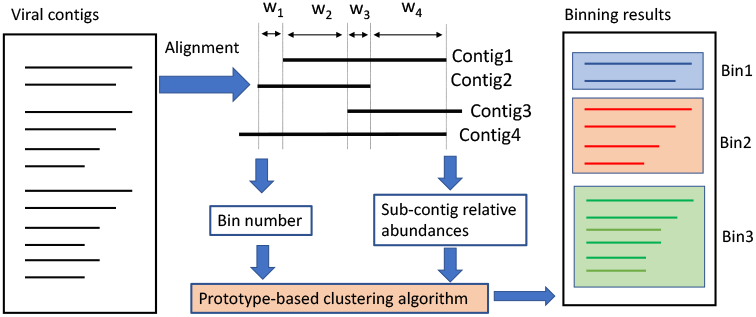
The pipeline of VirBin.

The underlying algorithm of grouping contigs into haplotypes is prototype-based clustering (Tan *et al.*, 2005). Features such as the overlaps and paired-end connections have limited usage in grouping distant contigs from the same haplotype. The clustering will mainly use the features based on the abundance distributions. Although abundance-based clustering has been used for contig binning from multiple samples (Wu *et al.*, 2014; Quince *et al.*, 2017), existing tools are not designed to tackle key challenges of distinguishing contigs of different haplotypes. First, the observed coverage of each contig not only depends on the abundance of the underlying haplotype, but also depends on whether it is a unique or shared region by two or more haplotypes. Second, heterogeneous coverage of each haplotype in an RNA viral quasispecies is common, which is caused by sequencing-related biases and compounded by gene expression. Thus, directly applying existing prototype-based clustering models such as Gaussian-mixture model to contigs is not expected to produce accurate clustering. Our solution to this problem is to cut the contigs into “windows” and to apply the clustering on sub-contigs that are more likely to represent one haplotype. In addition, instead of assuming any parametric distribution, which is usually not the case for haplotype contigs, we will use a non-parametric distribution.

### 2.1 Step 1: estimate the number of haplotypes via contig alignment

Although the high similarity between haplotypes presents a barrier to adoption of kmer-based features for distinguishing contigs from different haplotypes, it brings opportunities for haplotype number estimation. With stringent alignment threshold, contigs that can be aligned with each other usually come from the same region of the virus and thus the number of aligned contigs can be carefully used for haplotype number estimation.

We progressively build multiple sequence alignments using contigs’ pairwise alignments. In this step, base-level accuracy of the alignment is not a major concern and thus progressive construction of the alignment between contigs can serve the purpose well. We first sort the contigs by their lengths in descending order. The longest contig will be used as the first reference. All the other contigs will be aligned to the reference using blast+ (Camacho *et al.*, 2009) to generate an alignment profile similar to multiple sequence alignment. Two types of alignments are kept from the output of blast+. One is the “glocal” alignment, which is local to the reference but global to the shorter contigs. The other is overlap alignment, which is the alignment between the suffix/prefix strings of the contigs. If not all the shorter contigs can be aligned to the reference contig, this process will continue by using the second longest contig as the reference until all the contigs are used. Fig. 2.(c) shows the alignment between contigs using the longest contig as the reference, which is usually produced for the most abundant haplotype.

**Fig. 2.**
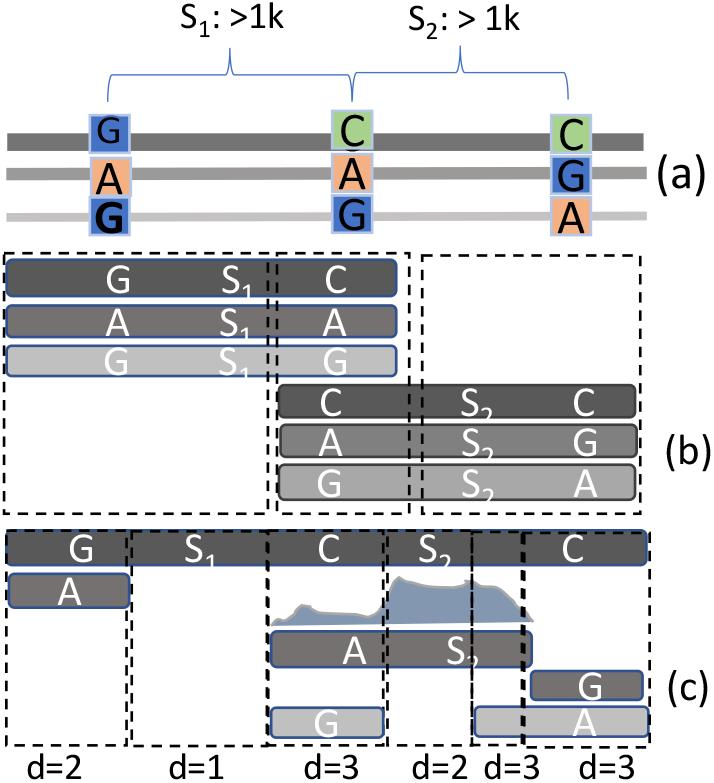
Window construction from aligned contigs. (a): Three haplotypes with mutations at three locations. Line weights represent the haplotype abundance. *S*_1_ and *S*_2_ are two mutation-free regions common to three haplotypes. *S*_1_ and *S*_2_ are at least 1k bp. (b): The alignment of contigs that satisfy the ideal condition. The grey-scale intensity represents the coverage of a contig. Three windows are produced. (c): The contigs that cannot cover all the three haplotypes. There are six windows. Their depth values are denoted below each window. For contig marked with “A *S*_2_ “, its sequencing coverage is plotted above the contig.

Each multiple alignment can be divided into many windows, which are formed whenever there is a change of the sequences in the alignment. We define the number of contigs inside each window as the *window depth, d*. Based on these definitions, we have the following observations.

When each position of the underlying haplotypes can be covered by at least one contig, *d* is equal to or larger than the number of haplotypes. Note that the common regions between different haplotypes are regarded as different positions and thus should be covered by different contigs. This conclusion can be proved by contradiction easily.

Fig. 2.(b) shows the contigs satisfying the conditions in the ideal case. There are three haplotypes with different abundances. They only contain mutations at three positions that are far away from each other. Because of the long common regions among them, assembly programs usually won’t be able to recover all the three genomes. Instead, they can generate short but correct contigs. In Fig. 2.(b), each position in the three haplotypes is covered by at least one contig. In this case, all the windows have depth of at least 3. If every position of a haplotype is only covered by one contig, the windows will have depth *N*. As some positions can be covered by multiple contigs, the overlaps between contigs contribute to window depth larger than *N*. For example, in Fig. 2.(b), the middle window contains the overlaps between two contigs from each haplotype and thus has depth 6. In this ideal case, we can choose the smallest window depth as the number of haplotypes in a sample.

In practice though, the assumptions about the contigs’ properties are not always true. Thus, in our implementation, we will use the consensus window depth as the number of haplotypes, by assuming that most windows cover all haplotypes and contain haplotype-specific mutations. For the contigs shown in Fig. 2.(c), window depth 3 is the most frequent one.

In the implementation, we first sort all the windows in descending order of window length. Then we choose the most frequent window depth of the top X windows as the number of haplotypes. The default value of X is 50 in our implementation. We will present the results of haplotype number estimation using consensus window depth for both simulated and real quasispecies data.

### 2.2 Step 2: contig clustering based on relative abundance distribution

Let the number of haplotypes estimated by step 1 be *N*. The problem can be defined as: given contigs *C*_0_, *C*_1_, …, *C_n_* assembled from viral quasispecies sequencing data, cluster the contigs into *N* groups so that each group contains contigs originating from the same haplotype. The relative haplotype abundance will be computed during the clustering process.

The clustering algorithm we adopt is prototype-based clustering and is essentially an augmented K-means algorithm. In a standard K-means algorithm, the centroid of the objects in a cluster is the prototype of the cluster. In our algorithm, the prototype is a distribution that is derived from the contigs and empirically describes the relative abundance distribution.

Unlike many existing contig clustering tools, our clustering is not applied to a complete contig. Because each contig can contain both haplotype-specific region and shared regions among different haplotypes, using the read coverage profile of the whole contig will confuse the clustering algorithm and makes the convergence slow or leads to wrong assignment of the objects. For example, in Fig. 2.(c), the contig “*A S*_2_ “ contains mutation *A* from one haplotype and also a shared region *S*_2_. Thus, significantly more reads will be mapped to the shared region and make the coverage for this contig highly heterogeneous. Thus, the objects as input to the clustering algorithm are “sub-contigs” in windows of depth *N*, where the sub-contigs are substrings of the contigs in these windows. They are more likely to represent the relative abundance of one haplotype.

The clustering algorithm will assign each sub-contig to one cluster based on the posterior probability of the abundance distribution. Although different clustering methods such as Gaussian mixture model can be applied to cluster the sub-contigs, the augmented K-means as shown below has the fastest convergence with better clustering accuracy according to our tests. Before we describe the main components, we first introduce the notations. The average relative abundance (denote as 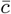) for a sub-contig *c_i_* in a window of depth *N* is calculated as:

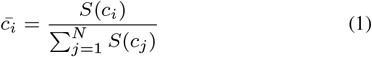

where *S* (*c_i_*) is the total number of reads covering sub-contig *c_i_*. Similarly, we can calculate the position-specific relative abundance vector 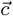 for a sub-contig *c_i_* as

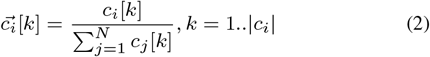

where *c_i_*[*k*] represents the reads coverage at position *k* of sub-contig *c_i_*. |*c_i_*| is the number of bases in the sub-contig.

*N* is the number of bins or haplotypes estimated by Step 1. VirBin utilizes the position-specific relative abundances of sub-contigs in windows with depth *N* to estimate the probability that a sub-contig belongs to a bin. Let the *N* bins be *H*_1_, *H*_2_, …, *H_N_*. Let *E_i_* be the abundance distribution for bin *i*. *E_i_*(*x*) is the probability of abundance variable *x* being generated by bin *i*.

The iterative clustering algorithm contains four steps as shown below:

#### Initialization

Initialize *N* groups by randomly assign sub-contigs to them.

#### Updating the abundance distribution E_i_

For each bin *i*, the component sub-contigs’ relative abundance profiles 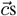 are aggregated to calculate the empirical probability density function *E_i_*. The aggregation is performed by calculating the normalized histograms for these relative abundance profiles, so that the summation of histogram values will be 1.

#### Re-assignment of the sub-contigs

Once *E_i_* is derived, the relative likelihood of *c_j_* being produced from the *i*th prototype distribution can be calculated as 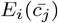. The prior probability of each bin (or haplotype) is a weighted sum of the likelihoods of all the component sub-contigs. The weights are determined by the total bases in the sub-contigs. The prior probability *Pr*(*H_i_*) for bin *i* is:

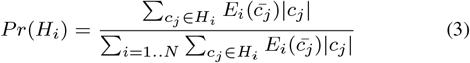

With both like lihood and prior from iteration *t*,the expected probability that *c_j_* belongs to haplotype *H_i_* at iteration *t* + 1 can be calculated as *likelihood* ⋆ *prior*, that is

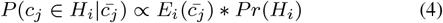

With the posterior probabilities calculated for each group distribution, we can reassign the sub-contig *c_i_* to the haplotype with the maximum posterior probability. The same reassigning procedures are applied for all the sub-contigs. With the assignment results, the distribution *E_i_* and prior probability *P* (*H_i_*) can be updated.

#### Iteration

Iterate step 2 and 3 until the clustering results do not change or the maximum number of runs have been achieved. The default maximum number of runs is 100.

### 2.3 Step 3: post-processing

The output of the augmented K-means is the clustered sub-contigs. For each cluster, its average abundance is calculated as the weighted average of the abundances of all sub-contigs in the cluster and the weight is determined by the length of asub-contig. The haplotypes’ abundances are the average abundances of the clusters.

As each contig can contain multiple sub-contigs, which could have different membership, the contig’s membership is determined by the dominant membership of its sub-contigs. For example, if a sub-contig is not in the window of depth *N*, it is not an input to the clustering step and will not be clustered. This could happen when a region of a contig is common to multiple haplotypes. It is also possible that the sub-contigs of a contig are assigned to different clusters, which could be caused by assembly errors.

## 3 Results

We evaluate the haplotype number estimation and clustering performance of VirBin on both simulated and mock HIV quasispecies sequencing data. The simulated data provide us with known ground-truth for accurate evaluation of the clustering performance. We produced simulated quasispecies sequencing data consisting of 5 haplotypes and 10 haplotypes, respectively.

For each experiment, we evaluate the performance of VirBin from three aspects: haplotype number estimation, clustering performance, and the computed haplotype abundance. When the originating haplotypes of the input contigs are known, we can evaluate both the recall and precision for the clustering step. First, we map the clusters to haplotypes based on the consensus haplotype label of the component contigs. If there is no consensus haplotype membership (e.g. a tie), we map the clusters to haplotypes based on the ranking of the abundance. Let a cluster be *B* and its paired haplotype be *H*. As the input to our program is a set of contigs, let the contig set originating from *H* be *C^H^*. Define *B* ∩ *C^H^* as the common regions between the two contig sets. Following other contig binning tools (Wu *et al.*, 2014; Lu *et al.*, 2017), the base-level recall for *H* is thus 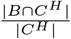, which quantified how many of the bases in *C^H^* are correctly clustered in *B*. The base-level precision is defined as 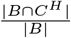, which quantifies how many of the bases in cluster *B* are from contig set *C^H^*. Similar metrics can be defined for contig-level, which can be found in the Supplementary Data file.

### 3.1 Simulated 5-haplotype quasispecies data

First, we constructed 5 haplotypes with high sequence similarity. Second, in order to simulate haplotypes of different relative abundances, we generated 3 sets of reads following different abundance distributions. Third, we generated contigs using two different methods. In method 1, we simulated 5 sets of error-free contigs of different sizes directly from the reference genomes. In method2, we applied available assembly tools to generate contigs from the reads. The simulated contigs are not dependent on any assembly tool and thus are ideal for evaluating the binning method. For simulated contigs, we have 3 (sets of reads) × 5 (sets of simulated contigs), i.e., 15 sets of input to our program. For assembled contigs, we have 3 (sets of reads) × 2 (sets of assembled contigs) as input to VirBin because we applied two assembly tools. Figure S1 in the Supplementary Data file sketches the process of input data generation. The data simulation details can be found below.

#### 3.1.1 Data simulation

##### HIV haplotype construction

There are many sequenced HIV strains in the HIV Sequence Database (Foley, Brian and Leitner, Thomas and Apetrei, Cristian, 2018). However, many of the strains do not possess sufficiently high similarity to be included in simulated quasispecies. Thus, we use both real and simulated strain sequences to simulate haplotypes of high similarity. Simulated strains were produced by mutating, deleting, or inserting bases at random positions from a real strain in the HIV database.

As a result, the five-haplotype dataset contains FJ061, 3 simulated haplotypes from FJ061, and FJ066. The sequence similarity between the simulated haplotype and its originating sequence is 97%. The average sequence similarity between all the five haplotypes is around 93%, which is comparable to the sequence similarity between haplotypes in a mock HIV quasispecies dataset (Jayasundara *et al.*, 2014).

##### Reads simulation using different haplotype abundance distributions

With available HIV haplotypes, simulated reads were generated from them by ART-illumina (Huang *et al*., 2012) as error-containing MiSeq paired-end reads, with read length of 250 bp, average insert size of 600 bp, and standard deviation of 150 bp. With the total coverage of 1000-x, three sets of reads are produced using different abundance distributions. The first one is based on the power law equation (Barbosa *et al*., 2012). The second and the third sets of reads represent challenging cases where different haplotypes have similar abundances, which create difficulties for abundance-based binning algorithms. The abundance differences in the second and third data set are 0.06 and 0.03, respectively. In total, there are 38,914 simulated reads for 5 HIV haplotypes. The relative abundance for five haplotypes in each read set can be found in Table 1. As the total coverage is 1000-x, the sequencing coverage of each haplotype is the product of the total coverage and the relative abundance.

**Table 1.** Relative abundance for 5 simulated HIV haplotypes in three read sets. “power” is the read sets generated based on the power law equation. “equal (6%)” is the abundance distribution with equal difference of 0.06. “equal (3%)” is the abundance distribution with equal difference of 0.03.

| Haplotype | Relative abundance % |  |  |  |  |
| --- | --- | --- | --- | --- | --- |
|  | FJ061 | FJ061-h1 | FJ061-h2 | FJ061-h3 | FJ066 |
| power | 39.0 | 23.3 | 16.0 | 12.8 | 9.0 |
| equal (6%) | 32.0 | 25.9 | 20.1 | 14.0 | 8.0 |
| equal (3%) | 26.0 | 23.0 | 20.0 | 17.0 | 14.0 |

##### Contig simulation

For each reference genome (denote its length as *L*), we randomly generated a list of location pairs (*p*_1_, *p*_2_), where 1 ≤ *p*_1_ < *p*_2_ ≤ *L*. Each location pair represents a candidate contig’s starting and ending position. Then, in the simulated contigs, we only keep the ones above 500 bp (i.e. *p*_2_ – *p*_1_ + 1 ≥ 500). In addition, we would like to simulate the hard case where the contigs cannot be extended any more using large overlaps. Thus, we sort all the remaining contigs by *p*_1_ and remove the ones that have overlaps of size above 100 bp with previous contigs in the sorted list. The five sets of simulated contigs have different N50 values and are referred to as “1000” to “5000”, indicating the upper bound of the contig length in each set. Table S1 in the Supplementary Data file shows the detailed properties of the five sets of contigs. All the simulated data sets can be downloaded from VirBin’s Github repository.

#### 3.1.2 Haplotype number estimation

According to our methods, the haplotype number estimation only depends on the alignment results of contigs. For all five sets of simulated contigs with different average lengths, the consensus window depth of the 50 longest windows is 5 for all. The histogram of window depth for 5 simulated contig sets is shown in Fig. 3. It is clear that window depth 5 dominates longest windows. Thus, the estimated number of haplotypes is 5, reflecting the truth for our data sets. In general, the percentage of windows with depth 5 increases with increasing contig lengths.

**Fig. 3.**
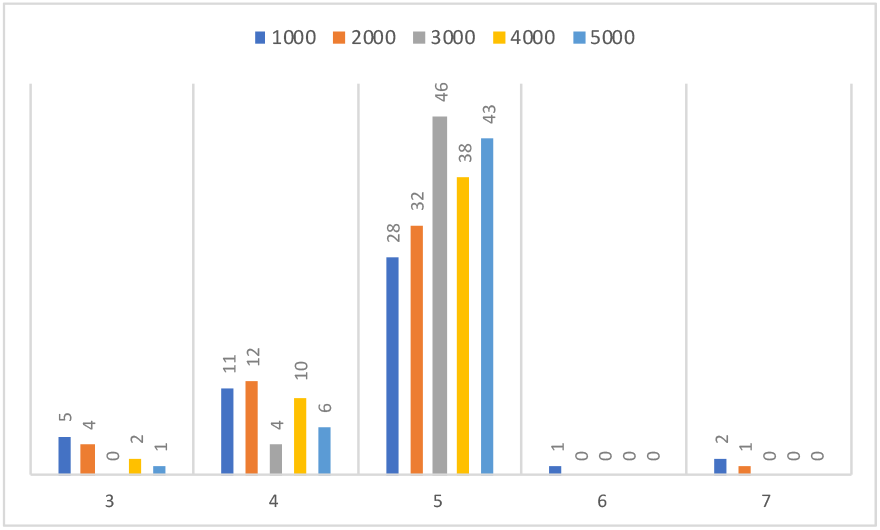
The histogram of window depth for the 50 longest windows. X-axis: the observed window depth, from 3 to 7. Y-axis: the number of windows with the corresponding window depth on X-axis. “1000” to “5000” represent the five sets of contigs.

#### 3.1.3 Clustering

We applied VirBin to cluster contigs into 5 groups. Since the ground truth about the haplotype membership of each contig is known, we were able to evaluate the clustering results by calculating the precision and recall at the base level. The evaluation results are shown in Fig. 4. The performance of clustering is worst for shortest contig set (denoted as 1000 along the Y-axis). With increasing contig lengths, the clustering performance becomes better for all three different abundance distributions. When the contigs are long, the clustering performance for haplotypes with different abundance distributions is comparable.

**Fig. 4.**
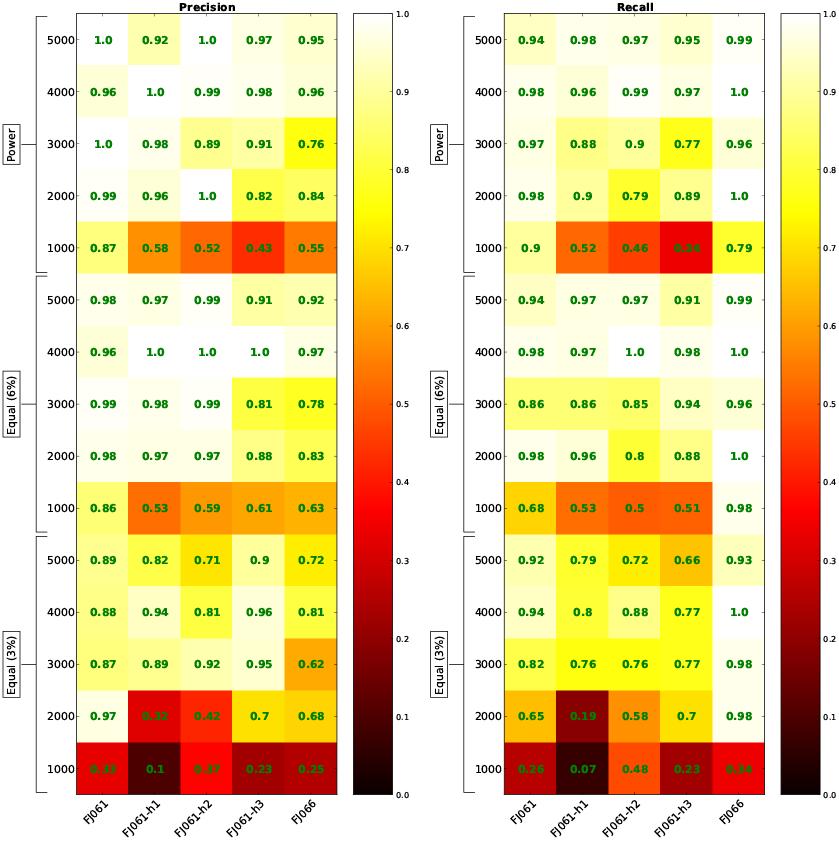
VirBin clustering results (recall and precision) on 15 simulated data sets. X-axis represents each haplotype, in decreasing order of relative abundance. Y-axis represents the 15 data sets.

The results were compared with MaxBin, which is a binning tool for metagenomic contigs based on tetranucleotide frequencies and reads coverage levels. MaxBin requires marker genes to identify seed contigs for binning. We were able to run the core clustering program of MaxBin by inputting both the number of haplotypes (i.e. 5) and the seed contigs manually. We randomly chose one contig from each haplotype as the seed contig and calculated the contigs’ abundances by mapping reads to them using Bowtie2. Although the haplotype number was explicitly provided to MaxBin, empty clusters can be produced by MaxBin. The results from MaxBin are shown in Fig. 5.

**Fig. 5.**
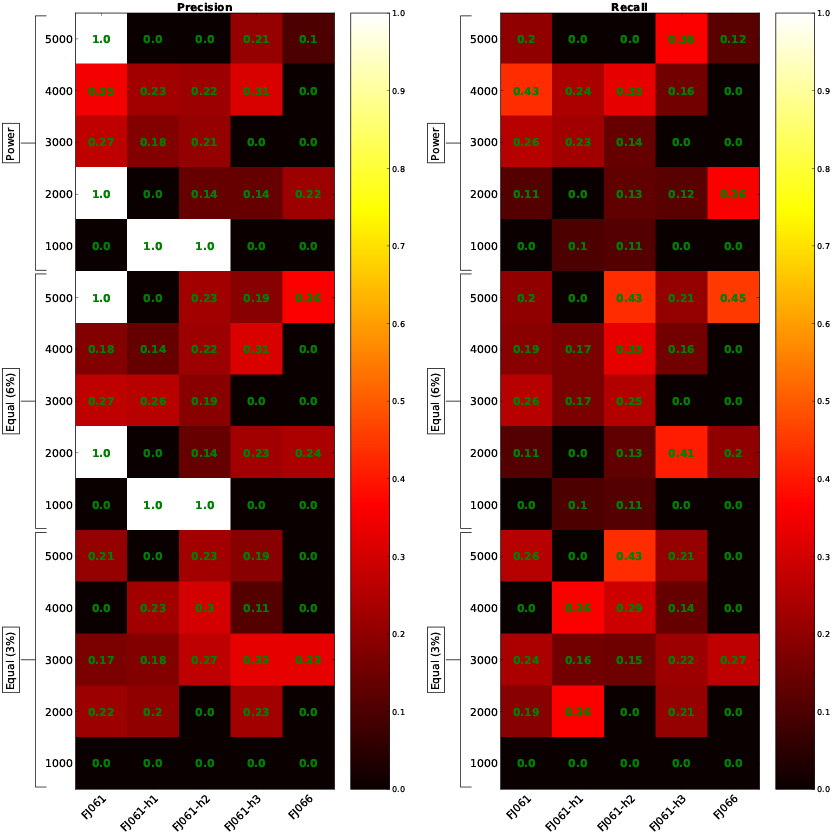
The recall and precision of contig binning results by MaxBin. X-axis represents each haplotype, in decreasing order of relative abundance. Y-axis is the index of the 15 data sets.

For the shortest contig set, MaxBin only reported two clusters with one contig for each cluster, leaving 59 (96.7%) contigs unclassified. For contigs sets from 2000 – 5000, MaxBin was able to generate five clusters, but with ~30% contigs unclassified. The results of MaxBin usually have lower precision or recall values than VirBin. In addition, for contig sets from 1000 – 4000, there are haplotypes without correctly assigned contigs. The lower sensitivity of MaxBin could be caused by its dependency on both sequence composition and contig coverage for clustering. Due to high sequence similarities between viral haplotypes, sequence composition is not informative enough in differentiating contigs from different viral strains. Instead, MaxBin could mistakenly cluster contigs from the homogeneous regions of the viral genome, leading to more chimeric clusters.

StrainPhlAn (Truong *et al.*, 2017) is also able to to characterize the genetic structure of viral strains in metagenomes. It takes the raw sequencing reads and MetaPhlAn2 (Truong *et al.*, 2015) database of species-specific reference sequences as input and aims to output the most abundant strain for each sample. However, it failed to detect any viral species at the first step running MetaPhlAn2. ConStrains (Luo *et al.*, 2015) is another tool designed to identify strain structures from metagenomic data. It uses bowtie2 to map reads to a set of universal genes and infers the within-species strains abundances by utilizing single-nucleotide polymorphism (SNP) patterns. This tool again did not get enough mapped reads to report any strain abundance. And it takes considerable efforts for us to modify their codes for our inputs. Thus, we cannot report the results from StrainPhlAn or ConStrains.

##### Relative abundance computation

Once the iterative clustering algorithm converges, the abundance of each cluster can be computed as the weighted average abundances for all contigs from this cluster.

Fig. 6 compares the known haplotype abundance profiles with our computed ones. There are three read sets with different abundance distributions (Table 1). For each distribution, there are five sets of contigs (Table S1 in the Supplementary Data file). Thus, three plots of five curves are presented to compare the ground truth and the computed abundance. In addition, we applied *χ*^2^-test to compare the ground-truth distribution and each computed abundance distribution. The p-values from all the tests are larger than 0.99, indicating that the distributions are not statistically different. As MaxBin only correctly clustered several contigs, we did not include the abundance comparison.

**Fig. 6.**
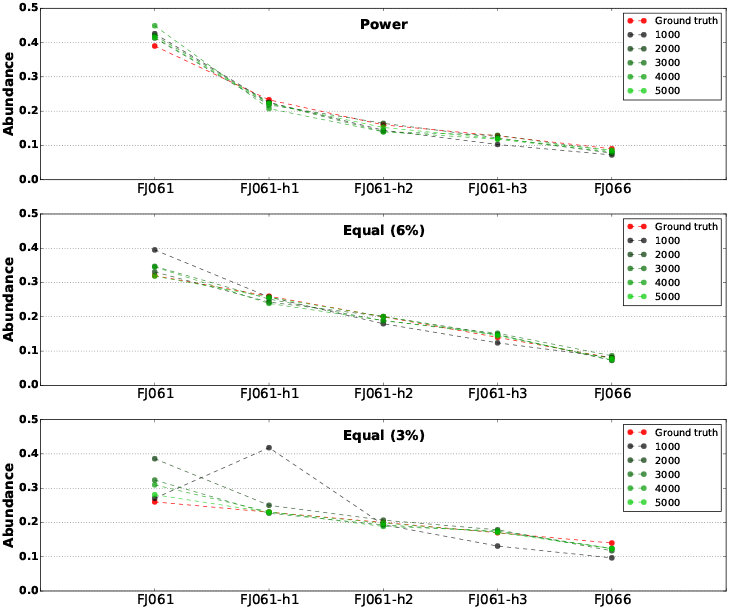
Compare the known haplotype abundance distributions with computed ones by VirBin. X-axis represents each haplotype. Y-axis is the ground truth or predicted abundance for each haplotype.

#### 3.1.4 Results on assembled contigs

In addition to simulated contigs, we also tested VirBin on assembled contigs by two *de novo* assembly tools: a generic assembly tool SGA (Simpson and Durbin, 2012) and a viral haplotype reconstruction tool PEHaplo. The assembled contigs were evaluated by MetaQuast (Mikheenko *et al.*, 2015) and the results are listed in Table S2 in the Supplementary Data file. Both PEHaplo and SGA produced enough contigs to cover almost all the five haplotypes. But contigs produced by PEHaplo have larger N50 values than contigs by SGA. The contigs are paired with haplotypes based on the highest sequence similarity. All of the contigs and their originating haplotypes have similarity of at least 98%.

For all three sets of contigs assembled by PEHaplo and SGA on three sets of reads, the consensus window depth of the 50 longest windows is 5, revealing the actual haplotype number.

Fig. 7.(a) presents the clustering results on contigs generated by PEHaplo and SGA. It shows that VirBin achieved good clustering results on contigs assembled by both assembly tools. The clustering results on SGA’s contigs are similar to PEHaplo’s contigs, with both high precision and recall. This observation is consistent with the results on simulated contigs that when the contigs are long enough, VirBin can produce good results.

**Fig. 7.**
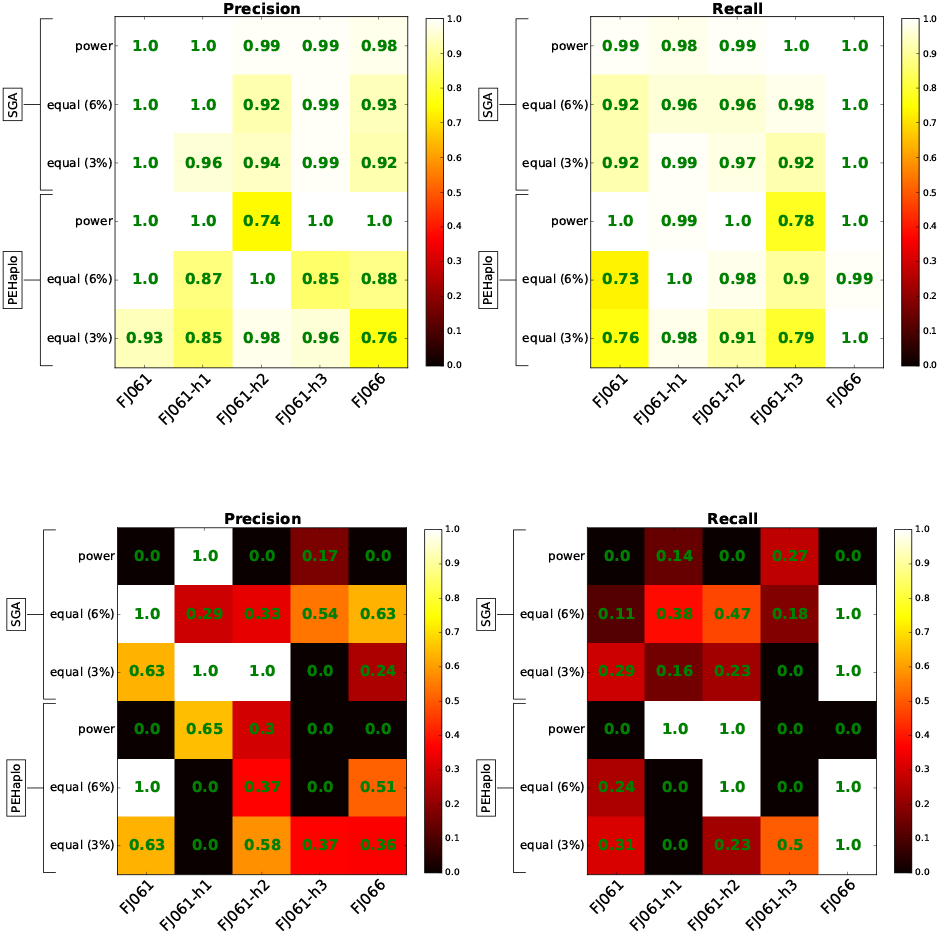
The clustering results of VirBin (a) and MaxBin (b) on contigs assembled by PEHaplo and SGA.

Again we compared our results with MaxBin. The clustering results of MaxBin on assembled contigs are shown in Fig. 7.(b). For contigs assembled by PEHaplo, MaxBin correctly clustered all corresponding contigs to the strain FJ061-h2 as the recall is 1.0. However, this cluster also involves many contigs from other strains as the precision value is low.

The comparison between the predicted abundance by VirBin and the ground-truth on two sets of assembled contigs is presented in Figure S2 in the Supplementary Data file.

We also simulated reads from 10 haplotypes and tested VirBin on this data set. The data generation and also the detailed results can be found in the Section 3 of the Supplementary Data file.

### 3.2 Mock HIV population MiSeq data set

In this experiment, we applied VirBin to a mock HIV quasispecies data set (SRR961514), sequenced from the mix of five HIV-1 strains (89.6,HXB2, JRCSF, NL43, YU2) with Illumina MiSeq sequencing technology (Di Giallonardo *et al.*, 2014). This data set contains 1,429,988 (250 bp) of reads that cover the five strains to 20,000x. The raw data set was preprocessed with FaQCs/1.3 (Lo and Chain, 2014) and Trimmomatic (Bolger *et al.*, 2014) to trim and filter low-quality reads or adapters. The remaining reads were then error-corrected with Karect (Allam *et al.*, 2015). After pre-processing, 774,044 reads were left. By mapping pre-processed reads to the available 5 reference genomes by bowtie2, we were able to estimate the abundance for each haplotype as shown in Fig. 8.

**Fig. 8.**
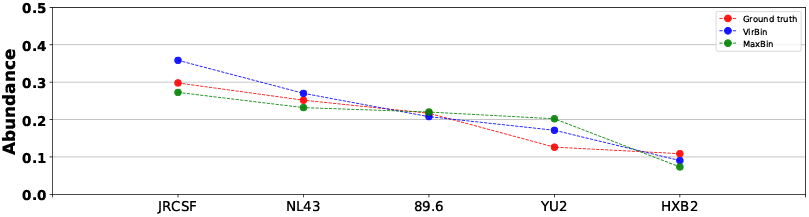
The true abundance distribution for real HIV quasispecies data and the computed relative abundance profiles by VirBin and MaxBin. The true average abundances sorted in descending order are: 29.79%, 25.18%, 21.68%, 12.62%, 10.87%.

We use the contigs assembled by PEHaplo as input for VirBin. PEHaplo produced 24 contigs from the real MiSeq HIV data set that can cover about 92% of the five HIV-1 strains. These contigs have a N50 value of 2,223 bp and the longest contig is 9,133 bp.

### Haplotype number estimation

VirBin was applied to the aligned contigs for haplotype number estimation. All the windows were sorted in descending order of window length. Out of the top 50 windows, 27 contain 5 contigs, 16 contain 6 contigs, and 2 contain 4 contigs. Out of the top 25 windows, 17 contain 5 contigs, 5 contain 6 contigs, and 1 contains 4 contigs. Similar to the simulated data, using the consensus window depth (i.e. 5) correctly predicted the haplotype number.

### Clustering results

The clustering algorithm was applied to cluster the contigs into 5 groups. For each contig, its originating haplotype is determined by comparing the contig with all reference genomes. The haplotype with the highest similarity and above 98% is assigned. The outputs of VirBin and MaxBin are shown in Table 2. StrainPhlAn and ConStrains were applied on this real HIV data set. StrainPhlAn was able to identify the HIV species, but could not report any strain information. ConStrains could not align enough reads to marker genes for further strain-level analysis.

**Table 2.** Base-level clustering results on assembled 5 real haplotype contigs for VirBin and MaxBin. The haplotypes are sorted in decreasing order of abundance.

|  | VirBin |  | MaxBin |  |
| --- | --- | --- | --- | --- |
|  | Precision (%) | Recall (%) | Precision (%) | Recall (%) |
| JRCSF | 65.1 | 50.8 | 0.0 | 0.0 |
| NL43 | 18.5 | 17.4 | 9.6 | 21.6 |
| 89.6 | 58.0 | 56.5 | 0.0 | 0.0 |
| YU2 | 34.0 | 30.0 | 56.6 | 50.9 |
| HXB2 | 48.5 | 70.2 | 39.1 | 27.0 |

Compared to the simulated contigs or assembled contigs using simulated reads, the results of VirBin on the real sequencing data have generally lower sensitivity and precision. There are two major reasons. First, the assembled contigs for real reads are more likely to contain errors. Second, this data set has several haplotypes with very similar average abundances. Referring to Fig. 8, the abundance difference between the 2 least abundant haplotypes is < 2%. Thus, the clustering algorithm could mix contigs from these haplotypes.

For the mock data experiment, we also present the recall and precision at contig level in Table S4 in the Supplementary Data file.

We again compared the predicted abundance profile with the known one in Figure 8. The *χ*^2^-test output p-value 0.9999 and 0.9995 for VirBin and MaxBin, respectively, indicating that the predicted abundance distributions by VirBin and MaxBin are not statistically different from the ground truth.

## 4 Discussion and conclusion

Overall, VirBin can cluster more contigs into their originating haplotypes than MaxBin. While VirBin focuses on sub-contigs that are more likely unique to one haplotype, MaxBin clusters whole contigs, which could contain regions common to multiple haplotypes and makes read coverage more heterogeneous. In addition, sequence composition-based features such as tetranucleotide frequencies are not effective in distinguishing highly similar viral strains.

Our experimental results show that VirBin works better for longer contigs with higher coverage of the underlying genomes. When the genome coverage by the contigs is below 70%, the performance of VirBin deteriorates because it becomes harder to estimate the correct number of haplotypes. In addition, the empirical experience shows that it is difficult to classify two viral strains when the abundance difference between them is below 3%. Thus, although we have demonstrated much better contig binning performance for distinguishing viral haplotypes than other contig binning tools, genome-scale viral haplotype construction is still a challenging problem.

## Funding

This work is supported partially by Michigan State University and City University of Hong Kong.

## Notes

https://github.com/chjiao/VirBin

